# Biomarker potential of real-world voice signals to predict abnormal blood glucose levels

**DOI:** 10.1101/2020.09.25.314096

**Authors:** Jouhyun Jeon, Adam Palanica, Sarah Sarabadani, Michael Lieberman, Yan Fossat

## Abstract

**Background:** Voice signal analysis is an emerging non-invasive technique to examine health conditions, and is implemented in various real-life applications and devices. The purpose of this study was to evaluate the association of voice signals with blood glucose levels in healthy individuals. The study aimed to investigate the longitudinal stabilities of voice signals and identify voice biomarkers to predict abnormal blood glucose levels.

**Methods:** We created voice profiles composed of 17,552,688 voice signals from 44 participants and their 1,454 voice recordings. From each voice recording, 12,082 voice-features were extracted. Longitudinal stabilities of voice-features were quantified using linear mixed-effect modelling. Voice-features that showed significant difference between different blood glucose levels, strong intra-stability and the ability to make distinct choice in decision trees were selected as voice biomarker. Voice biomarkers were fed into a multi-class random forest classifier to predict high, normal, and low blood glucose levels.

**Findings:** In total, 196 voice biomarkers were characterized. Results showed a predictive model with an overall accuracy of 78.66%, overall AUC of 0.83 (95% confidence interval is 0.80 – 0.85), and 0.41 of Matthews Correlation Coefficient (MCC) to discriminate three different blood glucose levels in an independent test set.

**Interpretation:** Our voice biomarkers could serve as a noninvasive and conventional surrogate of blood glucose monitoring in daily life as well as a screening tool to estimate potential risk of poor glycemic control.

**Funding:** This research was internally funded and received no specific grant from any funding agency in the public, commercial, or not-for-profit sectors.

## Introduction

Human voice is composed of complex signals that are tightly associated with physiological changes in body systems^1^. Due to the depth of signals that can be analyzed, as well as the wide range of potential physiological dysfunction that manifest in voice signals, voice has quickly gained traction in healthcare and medical research. For example, it has been shown that thyroid hormone imbalance caused the hoarseness of voice, and affected larynx development^2^. Unstable pitch and loudness were observed in patients with multiple sclerosis^3^. Other recent studies also demonstrated distinct voice characteristics that were associated with various pathological, neurological, and psychiatric disorders, such as congestive heart failure^4^, Parkinson’s disease^5^, Alzheimer’s disease^6^, post-traumatic stress disorder^7^, and autism spectrum disorder^8^. The human voice is now considered as an emerging biomarker, which is inherently non-invasive, low-cost, accessible, and easy to monitor health conditions in various real-life settings.

Glucose is an essential component of cellular metabolism, and its concentration in blood is regulated and maintained in a controlled, physiological range as a part of metabolic homeostasis^9^. Long-lasting disturbances in blood glucose concentrations can cause diabetes and diabetes-related complications. Diabetes has a high incidence (10.5% of population in 2018) and is one of the main causes of death in the United States (7^th^ leading cause)^10^. In spite of such risks, screening undiagnosed patients is not conducted routinely, and thus about 50% of adult diabetes cases are estimated to be undiagnosed, globally^11^. Therefore, implementing a regular monitoring tool for blood glucose into real-life is essential and urgent to identify potential individuals at a high-risk of developing prediabetes or diabetes when they are still healthy or asymptomatic. Voice analysis in healthy individuals can provide an efficient preliminary evaluation to examine glycemic control, which would be vital to reduce the medical and economic burden of diabetes. Since blood glucose levels can directly impact the neurologic, vascular, and muscular systems^12–14^, all of which are essential components of voice function, it is reasonable to assume that changes in the level of blood glucose can subtly influence aspects of voice even in healthy individuals. Recent studies have investigated whether Type 2 Diabetes patients have different voice characteristics compared to healthy controls^15,16^, and a higher vocal pitch has been observed as a potential clinical symptom of hypoglycemia in Type 1 Diabetes patients^17^. However, voice characteristics associated with abnormal blood glucose levels (*e.g*., elevated blood glucose not considered clinically hyperglycemic) in healthy or potentially prediabetic individuals remains unknown despite their considerable potential for clinical diagnostic utility.

Here, we investigated whether blood glucose levels were manifested in the voice of healthy individuals. To do this, we measured blood glucose levels of individual participants in an uncontrolled setting as they went about their daily lives, and had them record voices using a typical smartphone at several times throughout the day. We then created voice profiles combining with various clinicopathological information, quantified the longitudinal stability of those profiles, and showed that voice biomarkers can directly reflect different levels of blood glucose. These data were used to create and validate a predictive model to classify high, normal, and low blood glucose levels in healthy individuals.

## Methods

### Study design and participants

For the study, 54 volunteers (aged ≥ 18 years) were recruited from Klick Inc., a technology, media, and research company in the healthcare sector based in Toronto, Canada. They were all employees of Klick Inc. and volunteered via the company’s intranet system. The study was performed in accordance with relevant guidelines and regulations, and informed consent was obtained from all participants prior to study entry. The study received full ethics approval from Advarra IRB Services (www.advarra.com/services/irb-services), an independent ethics committee. Participants’ blood glucose levels were measured using a FreeStyle Libre glucose monitoring device (Abbott Diabetes Care), and voice samples of simple spoken sentences (e.g., “Hello, how are you? Today is September 5, 2019, 04:06 pm”) were recorded using participants’ smartphones. After the 14 days of collection of blood glucose levels and voice samples, data from seven participants were eliminated because of a malfunctioning glucose monitoring device (*e.g*., erroneous or missing measurements), and from one participant who failed to record proper voice sample. In total, 44 participants, and their 1,454 voice recordings with matched blood glucose levels were selected and used for further analyses. From the each voice recording, 12,072 voice-features were extracted using OpenSmile software (v.2.3.0), an open-source audio feature extractor^18^. The profiles of 17,552,688 voice signals (1,454 recording × 12,072 voice-features) were finally generated. Profiles was divided into two groups, Group A and Group B. Group A (1,290 voice recording from 39 participants) was used to extract features, measure intra-stability, identify voice biomarkers, and train a predictive model. Group B (164 voice recording from 5 participants) was used as an independent test set to evaluate a predictive model.

### Intra- and inter variance quantification and generalized intra-stability estimation of voice-features

The relative effects of intra- and inter-variance derived from participants as well as high, normal, and low blood glucose (BG) groups were assessed via linear mixed-effects modelling using the lme4 package (v1.1-21) in R statistical environment. In the model, BG groups and participants were specified as random factors to control for their associated intra-class correlation,

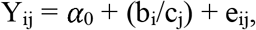

where Y_ij_ represents values of BG group *i* in participant *j*, *α*_0_ is a constant, b_i_ and c_j_ are the random effects for BG group *i* and participant *j*, respectively. Intercept varies among BG groups and participants within a BG group (expressed as b_i_/c_j_). e_ij_ is an unknown vector of random errors. To estimate generalized intra-stability, we calculated the intraclass correlation coefficient (ICC):

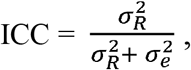

Where R represents random effects, b and c (b/c). The ICC represented the proportion of inter- b/c variance relative to total intra- and inter- b/c variance explained by a model. A high ICC indicates high generalized intra-stability within a BG group and participants within a BG group. ICCs of voice-features were estimated using Group A participants.

### Predictive model generation

To generate a predictive model that distinguishes abnormal high and low BG groups from normal BG group, 196 voice biomarkers were identified, and fed into a multi-class random forest (RF) classifier. The training set (Group A) and the RandomForestClassifier function built in the sklearn package (v.0.23.2) was used to train a model. To find optimal RF parameters (n_estimator, max_depth, max_features, and class_weight), grid search with 5-fold crossvalidation was conducted. Five-fold cross-validation set was generated using a stratified group K-fold method so that each fold has the same ratio of high, normal and low BG groups. Optimal parameters were determined based on the rank product of balanced accuracy (BCC), overall accuracy (ACC) and Matthews correlation coefficient (MCC). Prediction performances (BCC, ACC, and MCC) were measured using the pycm package (v.2.8) and sklearn package (v.0.23.2). Final model was trained on an entire training set with optimal parameters. To achieve the generalizability of a predictive model, we repeated this procedure five times. In each repeat, a cross-validation set was composed of different participant samples but kept the same BG group ratio. Finally, the ensemble model was built by combining all the results from five RF classifiers. The ensemble model was applied to an independent test set (Group B). Multi-class ROC was measured using the multiROC library (v.1.1.1) in R.

### Statistical analysis

Linear-mixed effect modelling and multi-class AUC estimation was performed using the programming language R (v3.4.0), and any remaining analyses were carried out in the programming language Python (v3.7.6) with the aforementioned packages. To examine the association of clinicopathological variables with blood glucose levels, p-values were measured using the Mann-Whitney U test for binary variables (sex and group), one-way ANOVA for multiple categorical variables (ethnicity), Spearman’s rank correlation coefficient for continuous variables (BMI, weight, height, diastolic blood pressure, and systolic blood pressure), and Kendall’s tau for ordinal variable (age group). A p-value of less than 0.05 was considered statistically significant. To evaluate the enriched audio-classes of voice-biomarkers, a hypergeometric test was performed. For the visualization of analyses, BPG library (v6.0.1) in R was used^19^.

## Results

### 1. Landscape of voice-features at different blood glucose levels

To understand the voice characteristics with respect to blood glucose (BG) levels, we collected 1,454 voice recordings at three different BG groups (70 low, 1,295 normal, and 89 high BG groups) from 44 healthy participants (**Figure 1A**) after the removal of unqualified voice recordings and participants (see **Supplementary Methods**). Participants were composed of 21 females and 23 males (**Figure 1B**). Study participants had an average age of 32 years and included various ethnic backgrounds (East Asian = 32%, Caucasian = 55%, South Asian = 2%, Middle Eastern = 2% and Other = 9%; **Table 1**). Clinicopathological variables (*e.g*., height, weight, blood pressure, and BMI) of participants were within the normal range (**Table 1**). For 14 days, each participant measured BG levels using a continuous glucose monitoring device (average BG level was 5.27 mmol/L). No statistically significant relationships between BG levels and clinicopathological variables were observed (p-value > 0.1; **Figure 1B**). On average, each participant provided 33 voice samples which were recorded at low (2 samples, BG level < 3.9 mmol/L), normal (29 samples, 3.9 mmol/L ≤ BG level ≤ 7.1 mmol/L), and high (2 samples, BG level > 7.1 mmol/L) BG levels across all time points (**Supplementary Figure 1**). Next, we divided our dataset into two groups. Group A (90% of the dataset, 1,290 voice recording from 39 participants) was used to characterize voice-features, evaluate their longitudinal stabilities, and build a predictive model to discriminate abnormal (high or low) BG levels from normal BG level. Group B (10% of the dataset, 164 voice recording from 5 participants) was used as an independent test set to evaluate the performance of the predictive model (**Figure 1A**).

**Figure 1.**
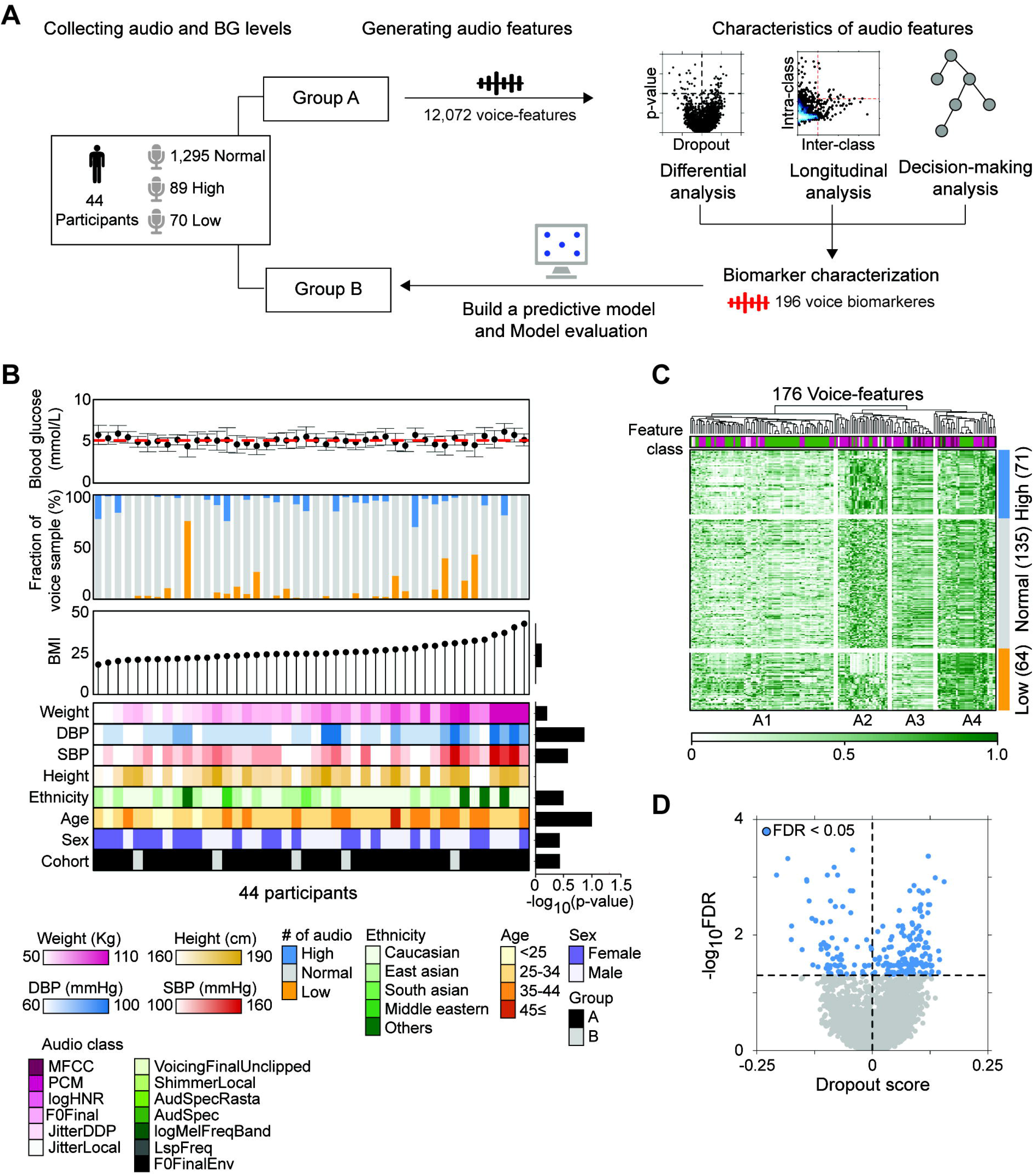
Voice profiles of healthy individuals. (A) Overview of analysis of voice signals and blood glucose (BG) levels in healthy individuals. (B) Landscape of BG levels, voice recordings, and clinicopathological information of 44 healthy individuals. Relationship between individual’s average BG levels and clinicopathological parameters were shown as p-values. (C) Profile of voice features. Values of 176 voice-features, which showed FDR < 0.05 and absolute dropout score > 0.05, were presented. (D) Volcano plot between dropout scores and FDRs of voicefeatures. Voice-features with FDR < 0.05 were coloured in blue.

**Table 1.**
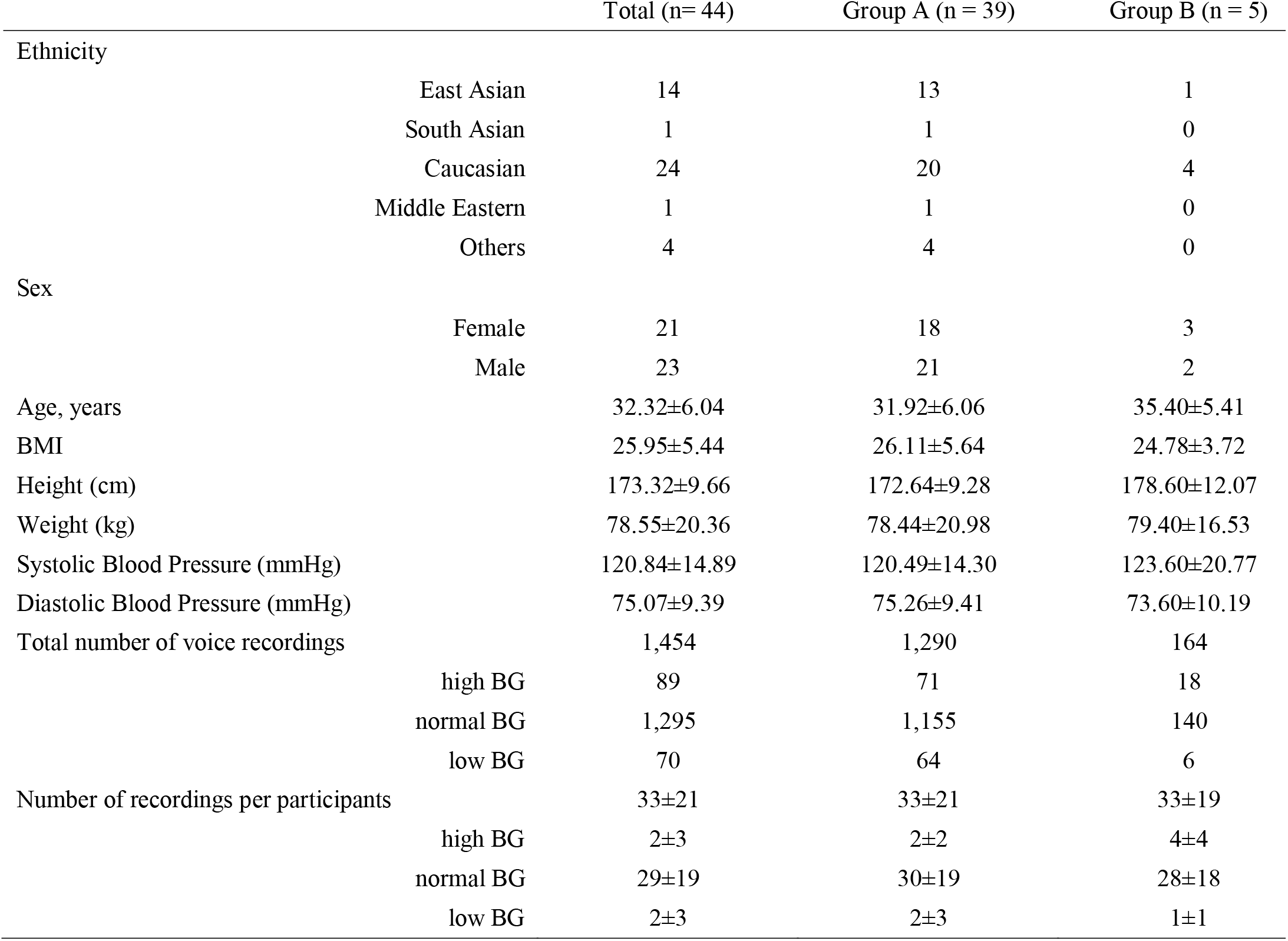
Demographics and audio characteristics

Voice-features at different BG groups were extracted and profiled from Group A participants. In total, 12,072 voice-features were identified using OpenSmile software^18^. These features represented 13 audio-classes representing different extractable signal components from a recorded voice (**Supplementary Table 1**). From the profile, we identified four clusters of voice-features (A1, A2, A3, and A4; **Figure 1C**). A2 and A3 showed the strongest signals in high BG level, and signals were reduced as BG levels decreased. They were mainly composed of Pulse-Code Modulation (PCM) and Mel-frequency cepstral coefficient (MFCC)-based features. Meanwhile, A1 and A4 showed reverse correlations between voice signals and BG levels and were mainly composed of sum of the auditory spectrum coefficients (AudSpec)-based features. Next, we investigated differences of feature signals among three BG groups (**Figure 1D**). To examine the directionality of signal changes, we measured dropout score (see **Supplementary Methods**). Negative dropout scores indicated the signal was increased as the BG level increased, whereas positive dropout scores indicated a signal that increased as the BG level decreased. The signals of 73 voice-features were significantly increased as the BG level increased (Dropout score < 0 and false discovery rate (FDR) < 0.05; **Figure 1D**). Of them, 42.47% were PCM-based features (**Supplementary Table 1**). Meanwhile, 153 features showed increased signals as BG levels were decreased (Dropout score > 0 and FDR < 0.05). Half of features (50.33%) were from AudSpec class.

### 2. Intra-stability of voice-features

To generate robust voice biomarkers, it is critical that voice signals remain stable over time within the same BG group and are distinctive between BG groups. To understand which voice-features were most and least stable within a BG group, we measured the between- and within-group variance of individual features and divided them into four quadrants (**Figure 2A**). We found that 106 voice-features were stable within a BG group (quadrant IV) showing high between-group variance (> top 1% of between-group variance) and low within-group variance (< bottom 99% of within-group variance). Meanwhile, another 106 voice-features were unstable within a BG group (quadrant II). Their within-group variances were more than 4 times as high as between-group variances. Over 98% (11,845) of voice-features showed nonsignificant between- and within-group variance (quadrant III), and 15 voice-features showed relatively high between- and within-group variances (quadrant I) implying that there could be additional factors that contribute to the stabilities of voice-features.

**Figure 2.**
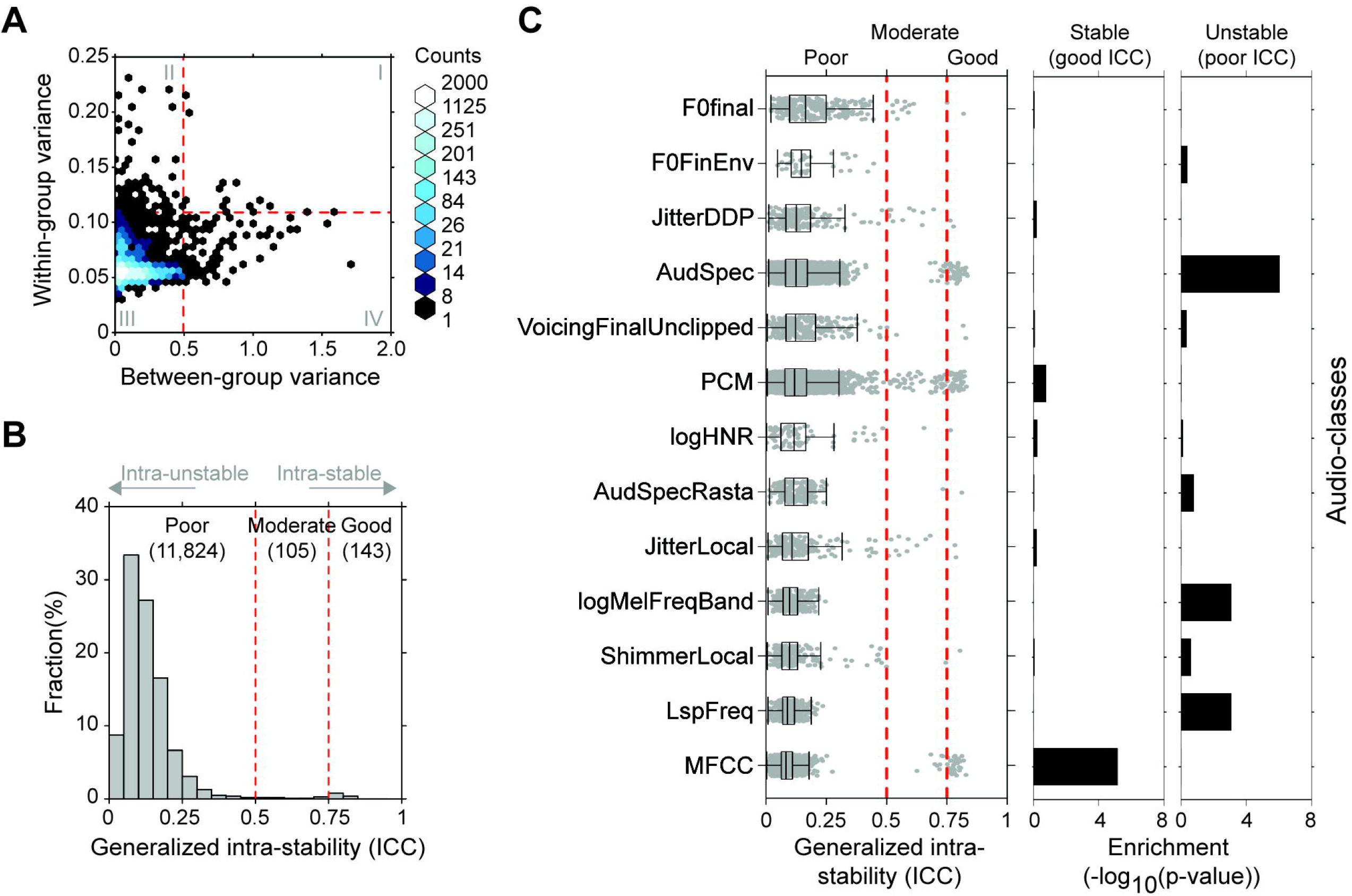
Intra-stability of voice-features. (A) Within- and between-BG group variance. Red dashed lines indicated top 1% of between-group variance (horizontal) and within-group variance (vertical). (B) The distribution of generalized intra-stability of 12,027 voice-features. Generalized intra-stability is estimated using intraclass correlation coefficient (ICC). (C) Distribution of ICCs depending on audio-classes. Enrichment of audio-classes in stable voice-features and unstable voice-features were shown.

Because of the potential to generate variations of voice signals within a participant resulting in increased variances within the same BG group, we decided to decode the variabilities derived from BG groups and participants, and estimated the generalized intra-stability of each voice-feature. To do this, we performed linear-mixed-effect modeling, and measured intra-class correlation-coefficient (ICC) as a metric for generalized intra-stability (**Figure 2B** and **2C**). The higher a voice-feature’s ICC, the more it is stable within a BG group across individuals. A majority of voice-features (11,824) showed a lack of stability within a BG group and participants within a BG group (unstable voice-features, poor ICC ≤ 0.5; **Figure 2B**), and 105 features showed moderate level of stability (0.5 < ICC ≤ 0.75). Only 143 (1.18%) voice-features were stable within a BG group across individuals (stable voice-features, ICC > 0.75). Interestingly, stable and unstable voice-features were enriched in different audio-classes (**Figure 2C**). Stable voice-features were significantly enriched in MFCC class (hypergeometric p-value, 7.03×10^−6^; **Figure 2C)**. Meanwhile, unstable voice-features were enriched in AudSpec (hypergeometric p-value, 9.27×10^−7^), logarithmic power of Mel-frequency bands (logMelFreqBand, p-value = 8.47×10^−4^) and line spectral pair frequency (LspFreq, p-value = 8.47×10^−4^) classes.

### 3. Voice-features associated with blood glucose levels

We then generated an optimal set of voice-features that could serve as biomarkers to discriminate between the three BG groups. We considered three criteria to select reliable biomarkers (**Figure 3A**). Features should show statistically significant differences between BG groups (traditional univariate analysis; *e.g*., small FDR), have high stability within the same BG group across participants (*e.g*., high ICC), and be relevant by having a sufficient ability to make a distinct choice in decision trees. To evaluate the decision ability of each voice-feature, we measured Gini impurity score and corrected them (Gini_c_) from multiple comparisons (see **Supplementary Methods**; **Figure 3B**). Gini impurity and Gini_c_ were positively related. Each voice-feature had 0.04±0.1 of Gini_c_ (0.08±0.13 of Gini impurity). 3,062 (25.36%) features were irrelevant (Gini_c_ = 0), and 4 features had significant abilities to make decisions on BG groups (Gini_c_ = 1). We selected 34 top ranked voice-features (Gini_c_ > 0.5), which were mainly composed of PCM (12 features), AudSpec (8 features), and MFCC (6 features) classes (**Supplementary Figure 2**).

**Figure 3.**
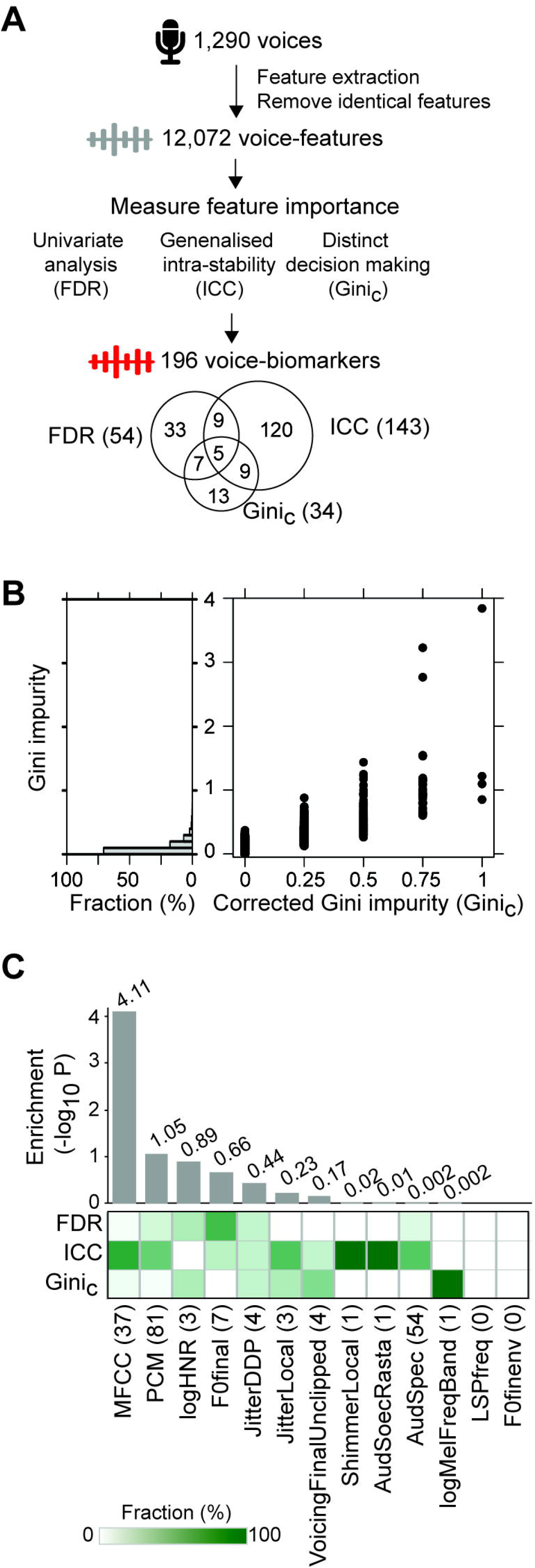
Identification of voice biomarkers. (A) Procedure to define voice biomarkers. In total, 196 voice-biomarkers were selected from three criteria (FDR, ICC, and Gini_c_). (B) Relevance of voice-features. Gini impurity scores were measured to evaluate the ability of each voice-feature to make distinct choice in decision trees (left), and were corrected from multiple comparisons (Gini_c_, right). (C) Enriched audio-classes of voice biomarkers. Hypergeometric p-values were shown on the top of bars.

In total, 196 voice-features were identified as a set of biomarkers (**Figure 3A**). They were composed of 33 FDR-specific (< 0.01), 120 ICC-specific (> 0.75), 13 Gini_c_-specific (> 0.5) features, and 30 biomarkers selected by at least two criteria. Biomarkers were involved in 11 out of 13 audio-classes (**Figure 3C**). The majority of biomarkers were involved in MFCC (37), PMC (81) and AudSpec (54) classes. The MFCC class was significantly enriched in our biomarkers set (p-value = 7.76×10^−5^). Furthermore, we found that biomarkers selected by different criteria were enriched in different audio-classes. For example, smoothed fundamental frequency contour (F0Fianl)-based biomarkers tended to be selected by FDR by having strong discriminatory power. MFCC-based biomarkers were likely to be selected by ICC indicating they were stable within a BG group and participants within a BG group. Voicing probability of the final fundamental frequency candidate with unclipped voicing threshold (VoicingFianlUclipped) and logMelFreqBand-based biomarkers were likely to be selected by Gini_c_ suggesting they had important roles to choose BG groups in decision trees. Taken together, our selected biomarkers could capture various profiles of the voice signals and avail information for the BG group classification.

### 4. Predictive models to classify distinct blood glucose levels

We then integrated our optimized voice biomarkers into a unified predictor that accurately discriminated between distinct BG groups (**Figure 4A**). Previously characterized 196 biomarkers were fed into a multi-class random forest (RF) classifier with hyperparameter optimization in the training set (Group A, 1,290 voice samples). We performed five-fold cross-validation to find an optimal set of parameters for a RF classifier and trained a predictive model (see **Methods**). To ensure generality of the prediction, we repeated the procedure five times by alternating voice samples in each fold and generated five different predictive models. Finally, the ensemble model was built by combining all the results from five models and applied to the independent test set (Group B, 164 voice samples). The ensemble model correctly predicted the BG groups in the test set (overall accuracy = 78.66%, balanced accuracy = 75.05%; **Table 2**). Over 80% of normal (recall = 80.71%) and low (recall = 83.33%) BG groups, and 61.11% of the high BG group were correctly predicted. The model had an overall Area Under the Curve (AUC) of 0.83 (micro AUC, 95% confidence interval (CI) = 0.80 to 0.85) and a corrected AUC of 0.71 (macro AUC, 95% CI = 0.64 - 0.77; **Figure 4B**). We also found that the predictive model outperformed any models generated by biomarkers which were selected by only FDR, only ICC and only Gini_c_. The predictive model showed the highest AUC (**Figure 4C**), and correctly predicted BG groups 1.07 ~ 2.53 times more than individual biomarkers selected by single or two criteria. Other performance measurements, Matthews Correlation Coefficient (MCC = 0.41) and corrected F1 score (macro F1 = 0.64), were 2.42±0.74 and 1.76±0.33 fold higher in the predictive model than single/double criteria-based biomarkers, respectively (**Table 2**). Additionally, to evaluate the null distribution of voice biomarkers, we generated 1,000 random sets of 196 voice-features and built a model from each. Indeed, our biomarker model outperformed the majority of random models across all performance evaluation metrics (**Figure 4D**).

**Figure 4.**
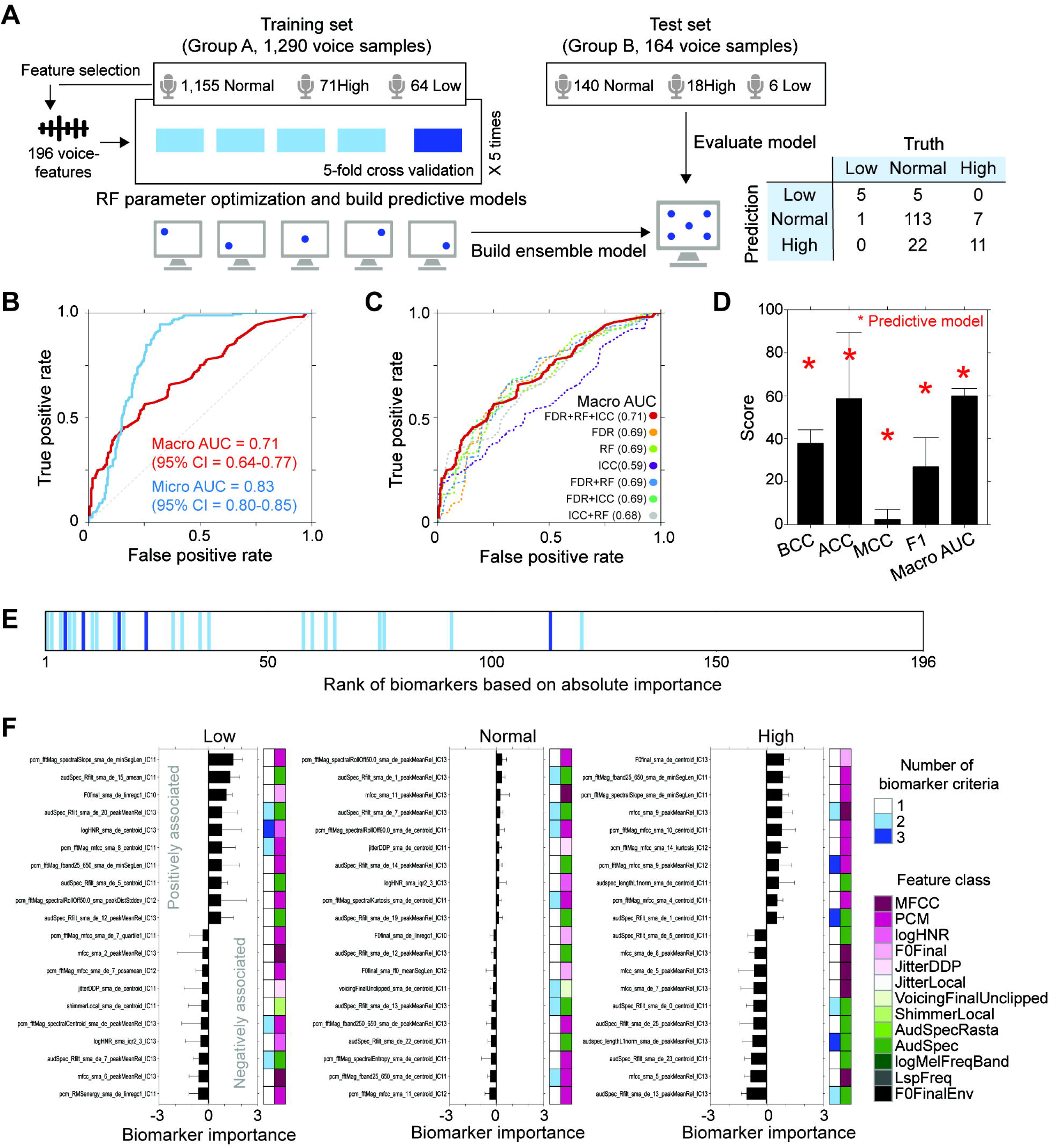
Evaluation of the predictive model. (A) Overview of the predictive model design. (B) The performance of the predictive model in the test set. Receiver operating characteristic (ROC) curves of micro average (blue) and macro average (red) were shown. (C) Performance of characterized voice biomarkers. Macro AUC of 196 biomarker-based predictive model (FDR+RF+ICC) was compared with those of models generated by individual biomarkers that were selected by only FDR, only RF, only ICC, FDR+RF, FDR+ICC, and ICC+RF. (D) Performance comparison between the predictive model and random models. Red asterisk indicated BCC, ACC, MCC, F1, and macro AUC of the predictive model. Error bars indicated standard deviation of performance matrix in 1,000 random models. (E) Importance of voice biomarkers to predict BG groups in the test set. Dard blue, light blue, and white bars indicated voice biomarkers that were selected by all three criteria, two criteria, and one criterion, respectively. (F) Relevant voice biomarkers to predict high, normal, and low BG groups. Top 10 voice biomarkers that were positively and negatively associated with BG groups were compared. Last four characters of voice-features (IC10, IC11, IC12, and IC13) indicated the origin of predefined feature set which OpenSmile provided.

**Table 2.** Performance of predictive models

| Features | BCC (%) | ACC (%) | MCC | Macro F1 | Macro AUC (95% CI) |
| --- | --- | --- | --- | --- | --- |
| FDR | 69.97 | 39.63 | 0.21 | 0.35 | 0.69 (0.64-0.72) |
| LMM | 52.17 | 39.02 | 0.13 | 0.33 | 0.59 (0.45-0.71) |
| Gini.c | 52.30 | 31.10 | 0.12 | 0.29 | 0.69 (0.64-0.73) |
| FDR + LMM | 59.18 | 65.24 | 0.22 | 0.48 | 0.69 (0.64-0.76) |
| FDR + Gini.c | 65.85 | 42.68 | 0.20 | 0.36 | 0.69 (0.64-0.74) |
| LMM + Gini.c | 61.53 | 49.39 | 0.20 | 0.45 | 0.68 (0.59-0.77) |
| FDR + LMM + Gini.c | **75.05** | **78.66** | **0.41** | **0.64** | **0.71 (0.64-0.77)** |
| Random* | 37.83 ± 6.28 | 58.74 ± 30.77 | 0.02 ± 0.05 | 0.27 ± 0.14 | 0.60 ± 0.03 |

\* 1,000 random models were generated using 196 randomly selected voice-features

Voice-biomarkers were selected from a training set using three criteria. To examine how much individual biomarkers contributed to the prediction of a test set, we performed Local Interpretable Model-agnostic Explanations (LIME) analysis, a technique to add interpretability and explainability to black box models^20^ and ranked 196 biomarkers based on their importance. We observed that biomarkers which were relevant in a training set also played important roles in predicting BG groups in the test set. Of 30 biomarkers selected by at least two criteria (**Figure 3A**), 20 (66.67%) were ranked within the top 50, and 28 (93.33%) were ranked within the top 100 relevant biomarkers to predict BG groups in a test set. Notably, 4 out of 5 (80%) biomarkers selected by all three criteria were ranked within the top 25 relevant biomarkers (**Figure 4E**). Next, we selected the top-10 positively and top-10 negatively associated biomarkers for BG group prediction to understand how biomarkers were combined and decided each BG group (**Figure 4F**). For the prediction of high BG level, PCM-based biomarkers were likely to be associated positively (*i.e*., high values affected correct prediction). Meanwhile, MFCC- and AudSpec-based biomarkers tended to be associated negatively with the prediction (*i.e*., low values affected correct prediction). For predicting low BG levels, AudSpec-based biomarkers were positively associated, showing their ability to track with both elevated and decreased BG level groups. In normal BG levels, jitter- and harmonic-to-noise ratio (HNR)-based biomarkers showed positive associations, which were opposite of their association for high BG prediction. AudSpec- and PCM-based biomarkers showed both positive and negative associations.

## Discussion

It has been shown that one-third of type 2 diabetes patients do not present symptoms until complications appear^21^, and undiagnosed diabetes is associated with higher risk of mortality compared to normoglycemic individuals^22^. Such diagnostic limitations suggested the need for effective screening techniques to differentiate an individual at high-risk from one at low-risk of having the disease in the future. Earlier identification of potential prediabetic individuals, and their monitoring and treatment can reduce the economic and social burden of diabetes and its complications. In this study, we demonstrated, for the first time, the association between voice signals and blood glucose levels in healthy individuals. Specifically, we identified 196 voice biomarkers to identify abnormally high and low BG levels. These voice biomarkers may serve as a non-invasive and conventional surrogate of blood glucose monitoring in daily life as well as a preliminary screening tool to identify individuals with potential prediabetes or those at risk of developing diabetes in the future.

We provided a new strategy to identify robust non-invasive voice biomarkers through parallel evaluation of feature importance. Repetitive voice recordings allowed us to quantify signal variances of voices within and between BG groups across all participants. From this longitudinal analysis, we could generalize intra-stabilities of voice-features and identify relevant biomarkers that present consistent signals to classify BG groups, regardless of time and individual to record voices. Traditional univariate analysis provided information to estimate the power of voice-features to discriminate abnormal BG groups. Lastly, Gini impurity score measured the probability of each voice-feature to decide a correct BG group in decision trees, and prioritized features. By integrating three biomarker selection strategies, we penetrated various different profiles of the voice-features and enhanced both accuracy and reliability of our predictive model.

Our biomarker discovery strategy successfully identified voice biomarkers that were physiologically associated with blood glucose levels and perhaps diabetes development. MFCC features have been studied to classify voices at risk for pathological conditions^23^ and to build a regression model to estimate blood glucose levels^24^. The other biomarkers, representing the changes of jitter, shimmer, loudness, and harmonic-to-noise ratio (HNR), captured the instability of oscillating patterns and closure of vocal folds. It has been shown that abnormal blood glucose levels caused the loss of fine motor muscle control^25^ and laryngeal sensory neuropathy^26^. Also, patients with Type 1 and 2 diabetes commonly showed dry mouth and decreased salivary flow rates^27^, which caused difficulty in phonation due to decreased lubrication mechanism of larynx^28^. Such physiological changes would affect vocal frequency and amplitude alternating phonation function.

In general, the normal hormonal changes in the morning increase blood glucose level regardless of health conditions to help individuals to have enough energy to get up and start the day^29^. Interestingly, voice sounds in the morning are relatively deeper compared to the sound during the day since vocal cords are relaxed (unused through night), swollen and thickened by the concentration of fluids in the upper body during sleeping. These unique physiological changes would affect the prediction of blood glucose levels from voices in the morning. Indeed, from our independent test set, we observed the lowest accuracy of BG level prediction in the morning between 6am to 12pm (25% of accuracy; **Supplementary Figure 3**). There were four voice samples that were recorded at high BG levels in the morning. Of them, three failed to predict BG levels correctly. Use of additional participants and their voice recordings may refine the assessment of longitudinal stability of voice features and improve biomarker discovery and time-dependent BG level prediction.

Overweight, high BMI, and high blood pressure are well known risk factors for both prediabetes and diabetes^30^. Integration of clinicopathological variables could improve the prediction accuracy of individuals, especially those at high-risk of disease in the future. Indeed, we observed that one individual in our test set (Group B) who had a relatively high BMI and blood pressure yielded low accuracy (42.85%) to predict BG groups. Meanwhile, four other healthy individuals, who showed a normal range of BMI and blood pressure, yielded 79.69% of accuracy to predicted BG groups (**Supplementary Figure 4**). We suspected that integration of abnormal range of clinicopathological signals may aid better prediction. One limitation of our study is that further investigation in larger and more diverse participants are required to ensure the generalizability of our findings.

Human voice signals can be a rich source of clinically relevant information while being non-invasive to measure, cost-effective, scalable, and accessible 24 hours a day in remote locations around the world. This work reinforces the idea that combining voice signals and machine learning techniques makes it possible to create a reliable and efficient system to identify abnormal blood glucose levels in otherwise healthy individuals. Glucose levels are traditionally measured with invasive continuous glucose monitoring (CGM) devices or finger prick tests. However, our novel method of analyzing voice biomarkers has the potential of being implemented in either healthy, prediabetes, or undiagnosed diabetes individuals during regular physician checkups. The fact that voice samples were also recorded on personal smartphones without any specific audio filters gives extra support for its potential use in everyday situations for patients of all demographics. The long-term implications include reducing specialized healthcare equipment costs and resources associated with diabetes-related treatment, as well as enhancing overall health and quality of life.

## Data sharing

The training and validation sets, which are composed of voice-features and blood glucose levels, will be available for academic purpose only after approval by corresponding author. The machine-learning algorithm is a third-party product that we have no rights to share.

## Acknowledgements

Thanks to all of the participants, project members, supporters, and researchers at Klick Inc. for the successful development, implementation, and evaluation of this research. We would like to thank Andrew Lee, Jeremy Jurksztowics, Anirudh Thommandram, and JJ Mifsud for technical supports and assistance. Also, we would like to thank Peter Leimbigler and Gaurav Baruah for supports in analysis. J.J., A.P., M.L., and Y.F. designed the study. A.P. and Y.F. collected the Data. J.J., and S.S. analyzed the data. J.J. interpreted the data, and wrote the manuscript with contributions from A.P. All authors revised the manuscript and contributed to the final review and editing, and have approved the final manuscript.

