## Supplementary Materials for "Biomarker potential of real-world voice signals to predict abnormal blood glucose levels"

### **Supplementary Methods**

#### ***Study population***

For the study, individuals who were below the age of 18 or those who were pregnant, or breastfeeding were excluded from the initial recruitment process. From the 54 volunteers, we further excluded two participants who were diagnosed with mental or physiological medical conditions and took prescription medication that could interfere with normal blood glucose regulation. The remaining 52 participants completed a self-report demographic survey, and had physiological variables measured, including height, weight, body mass index (BMI), systolic blood pressure, and diastolic blood pressure.

#### ***Measuring blood glucose levels***

To measure blood glucose levels, the FreeStyle Libre glucose monitoring device (Abbott Diabetes Care; https://myfreestyle.ca/en/products/libre) was used. It measured blood glucose levels (in mmol/L) at 15-minute intervals with a minimally invasive 5 mm flexible

filament inserted into the posterior upper arm. The device provided consistent accuracy and reliability throughout the 14 days regardless of age, sex, body weight, BMI, or time of use (day versus night)^1,2^. Measured blood glucose (BG) levels were divided into three BG groups based on general blood glucose level for non-diabetic individuals^3^. High BG indicated elevated BG levels (BG level > 7.1 mmol/L), and low BG indicated reduced BG levels (BG level < 3.9 mmol/L) compared to the normal range of BG levels (normal BG, 3.9 mmol/L ≤ BG level ≤ 7.1 mmol/L).

#### ***Collecting and pre-processing voice samples***

A custom mobile software application was built by Klick Inc. to record voice samples using participants’ smartphones (iOS and Android compatible). The downloaded app required users to input a unique participant identification code provided to them at study initiation, and then allowed them to make voice recordings using their own smartphone. All recordings were timestamped and immediately uploaded to a secure cloud storage system, accessible only to researchers. Throughout the entire study period (14 continuous days), participants were asked to record their voice via their smartphone at least 5 random times (of their choice) throughout the day, with the following phrase: “Hello, how are you? Today is [current day’s month, day, year, and time]”. During recordings, the mobile app displayed the specific reading instructions for the exact sentence to speak (*e.g.*, Read: “Hello, how are you? Today is September 5, 2019, 04:06 pm”). The app would immediately update the new reading instruction based on the relevant date and time.

Next, to maintain high quality recordings, voices that were recorded with partial sentences, unknown words, excessive background noise, and multiple voices (e.g., others speaking in the background) were excluded (363 recordings). To increase the volume of digital audio and have appropriate sample amplitude range, all voice recordings were normalized. Then, dynamic compression was performed to get audibility for low-level passages without reaching uncomfortable loudness levels for high-level signals^4^. Voice recordings were re-normalized after dynamic compression. Next, only active human voices were extracted using voice activity detection (VAD) techniques. These audio preprocessing were performed using python package webrtcvad (v.2.0.10) and SoX software (v. 14.4.2). After the pre-processing, 1,454 voice recordings from 44 participants were mapped to corresponding blood glucose levels, which were the nearest measurement from a given voice recording (within ± 15 minutes) and used for analyses.

#### ***Voice-feature extraction and profiling***

To extract and profile voice-features, we employed OpenSmile software (v.2.3.0), an open-source audio feature extractor^5^. It united feature extraction algorithms that represented 13 different aspects (classes) of voice signal and phonatory function : (1) Mel-frequency cepstral coefficient (MFCC), (2) logarithmic harmonic-to-noise ratio (logHNR), (3) smoothed fundamental frequency contour (F0Final), (4) envelope of smoothed F0Final (F0FinalEnv), (5) difference of period lengths (JitterLocal), (6) difference of JitterLocal (JitterDDP), (7) voicing probability of the final fundamental frequency candidate with uncliped voicing threshold (VoicingFianlUnclipped), (8) amplitude variations (ShimmerLocal), (9) sum of the auditory spectrum coefficients (AudSpec), (10) relative spectral transform of AudSpec (AudSpecRasta), (11) logarithmic power of Mel-frequency bands (logMelFreqBand), and (12) line spectral pair frequency (LspFreq), and (13) pulse-code modulation (PCM) that extract spectral features such as spectral energy, roll-off, flux, centroid, entropy, variance, skewness, kurtosis, sharpness, and loudness. Four pre-defined feature sets that OpenSmile provided were used to extract voice-features. They were composed of features that were used for Interspeech 2010 paralinguistic Challenge (IC10), Interspeech 2011 speaker state Challenge (IC11), Interspeech 2012 speaker trait Challenge (IC12), and Interspeech 2013 ComParE Challenge (IC13). In total, we extracted 12,072 voice-features after the removal of identical feature values. All feature values were re-scaled to have values ranging from 0 to 1:

Re-scaled feature value = $\frac{(V_{\mathrm{ij}} - \mathrm{Min}_{i})}{(\mathrm{Max}_{i} - \mathrm{Min}_{i})}$ ,

where V_ij_ indicated a value of feature *i* in sample *j*. Min_i_ and Max_i_ represented the minimum and maximum value of feature *i* in all samples, respectively.

#### ***Measuring the association between voice signals and blood glucose groups***

To incorporate voice signals from multiple time points in a profile, a dropout score was introduced. Dropout score assigned a value of each voice-feature by calculating the difference between feature value at each BG group and the value at the high BG group.

Dropout score = $\frac{1}{2}\times$ ((N_i_ - H_i_) + (L_i_ - H_i_)),

where Hi, Ni and Li are average values of feature *i* in high, normal and low BG groups, respectively. Positive dropout score indicated feature values were increased as the BG level decreased (H_i_ < N_i_ < L_i_). Negative dropout score indicated feature values were increased as the BG level increased (H_i_ > N_i_ > L_i_).

#### ***Biomarker characterization***

The selection of reliable voice biomarkers reduces the dimensionality of the feature space, avoid overfitting, and achieves better generalizability. Voice biomarkers were defined using three criteria. First, voice biomarkers showed significantly different values between BG groups. One-way analysis of variance (ANOVA) was used to examine statistical differences, and Benjamini-Hochberg-adjusted P-values were used to account for multiple-comparisons testing. Biomarkers showing p-values < 0.01 were selected. Second, voice biomarkers showed intra-stability within a BG group and participants within a BG group. Voice-features showing ICC > 0.75 were defined as biomarkers. ICC cutoffs 0.5 and 0.75 indicated good and moderate reliability, respectively^6^. Lastly, voice biomarkers should have sufficient ability to make distinct predictions in decision trees. To evaluate the decision ability of voice-features, Gini impurity scores were measured using the RandomForestClassifier function built in the sklearn package (v.0.23.2) in Python. Gini impurity scores were corrected through 1,000 repeated random stratified subsampling to generalize feature relevance. For each iteration, Gini impurity scores were measured from the randomly selected 29 participants in Group A, and scores were normalized to have a same range of values (normalized Gini impurity score, Gini_n_):

Gini_ni_ = $\frac{{Gini impurity}_{i}}{\sigma}$

Where, Gini impurity_i_ indicates Gini impurity score of voice-feature *i*, μ and σ indicate mean and standard deviation of Gini impurity scores. Each voice-feature has 1,000 Gini_n_, and finally we measured corrected Gini impurity score (Gini_c_):

Gini_c_ = 1 - $\frac{n}{1000}$

Where n indicated the number of Gini_n_ whose absolute value ≥ 1.96. Biomarkers are defined when they have Gini_c_ > 0.5. In total, 196 voice-features were defined as voice biomarkers and fed into a predictive model to identify distinct BG groups.

#### ***Interpretation of the predictive model***

To understand how each voice biomarker contributed to the prediction of a test set, Local Interpretable Model-agnostic Explanations (LIME) analysis was performed^7^. Lime provides three types of weights per voice biomarker. Each weight represented the contribution to predict high, normal and low BG groups in a given sample. To evaluate the importance of voice biomarkers in a high BG group, we only compiled high BG weights from voice samples predicted as a high BG group, and ranked voice biomarkers based on their average weight. Importance for normal and low BG groups also followed the same procedure. lime package (v.0.1) in Python was used for analyses.

#### ***Statistical analysis***

Linear-mixed effect modelling and multi-class AUC estimation was performed using the programming language R (v3.4.0), and any remaining analyses were carried out in the programming language Python (v3.7.6) with the aforementioned packages. To examine the association of clinicopathological variables with blood glucose levels, p-values were measured using the Mann-Whitney U test for binary variables (sex and group), one-way ANOVA for multiple categorical variables (ethnicity), Spearman’s rank correlation coefficient for continuous variables (BMI, weight, height, diastolic blood pressure, and systolic blood pressure), and Kendall’s tau for ordinal variable (age group). A p-value of less than 0.05 was considered statistically significant. To evaluate the enriched audio-classes of voice-biomarkers, a hypergeometric test was performed. For the visualization of analyses, BPG library (v6.0.1) in R was used^8^.

#### ***Supplementary References***

1	Hoss U, Budiman ES, Liu H, Christiansen MP. Continuous glucose monitoring in the subcutaneous tissue over a 14-day sensor wear period. *J Diabetes Sci Technol* 2013. DOI:10.1177/193229681300700511.

2	Bailey T, Bode BW, Christiansen MP, Klaff LJ, Alva S. The Performance and Usability of a Factory-Calibrated Flash Glucose Monitoring System. *Diabetes Technol Ther* 2015. DOI:10.1089/dia.2014.0378.

3	Alvi GB, Qadir MI, Ali B. Assessment of Inter-Connection between Suriphobia and Individual’s Blood Glucose Level: A Questionnaire Centred Project. *J Clin Exp Immunol* 2019; **4**.

4	Kirchberger M, Russo FA. Dynamic Range Across Music Genres and the Perception of Dynamic Compression in Hearing-Impaired Listeners. In: Trends in Hearing. 2016. DOI:10.1177/2331216516630549.

5	Eyben F, Wöllmer M, Schuller BB, Weninger F, Wollmer M, Schuller BB. OPENSMILE: open-Source Media Interpretation by Large feature-space Extraction. *MM’10 - Proc ACM Multimed 2010 Int Conf* 2015. DOI:10.1145/1873951.1874246.

6	Koo TK, Li MY. A Guideline of Selecting and Reporting Intraclass Correlation Coefficients for Reliability Research. *J Chiropr Med* 2016. DOI:10.1016/j.jcm.2016.02.012.

7	Ribeiro MT, Singh S, Guestrin C. ‘Why should i trust you?’ Explaining the predictions of any classifier. In: Proceedings of the ACM SIGKDD International Conference on Knowledge Discovery and Data Mining. 2016. DOI:10.1145/2939672.2939778.

8	P’ng C, Green J, Chong LC, *et al.* BPG: Seamless, automated and interactive visualization of scientific data. *BMC Bioinformatics* 2019. DOI:10.1186/s12859-019-2610-2.

### **Supplementary Figures**

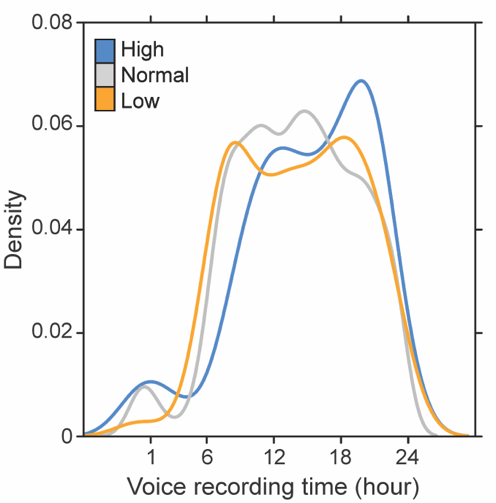

**Supplementary Figure 1.** Distributions of voice recording times. Blue, grey, and yellow indicated high, normal, and low blood glucose levels, respectively.

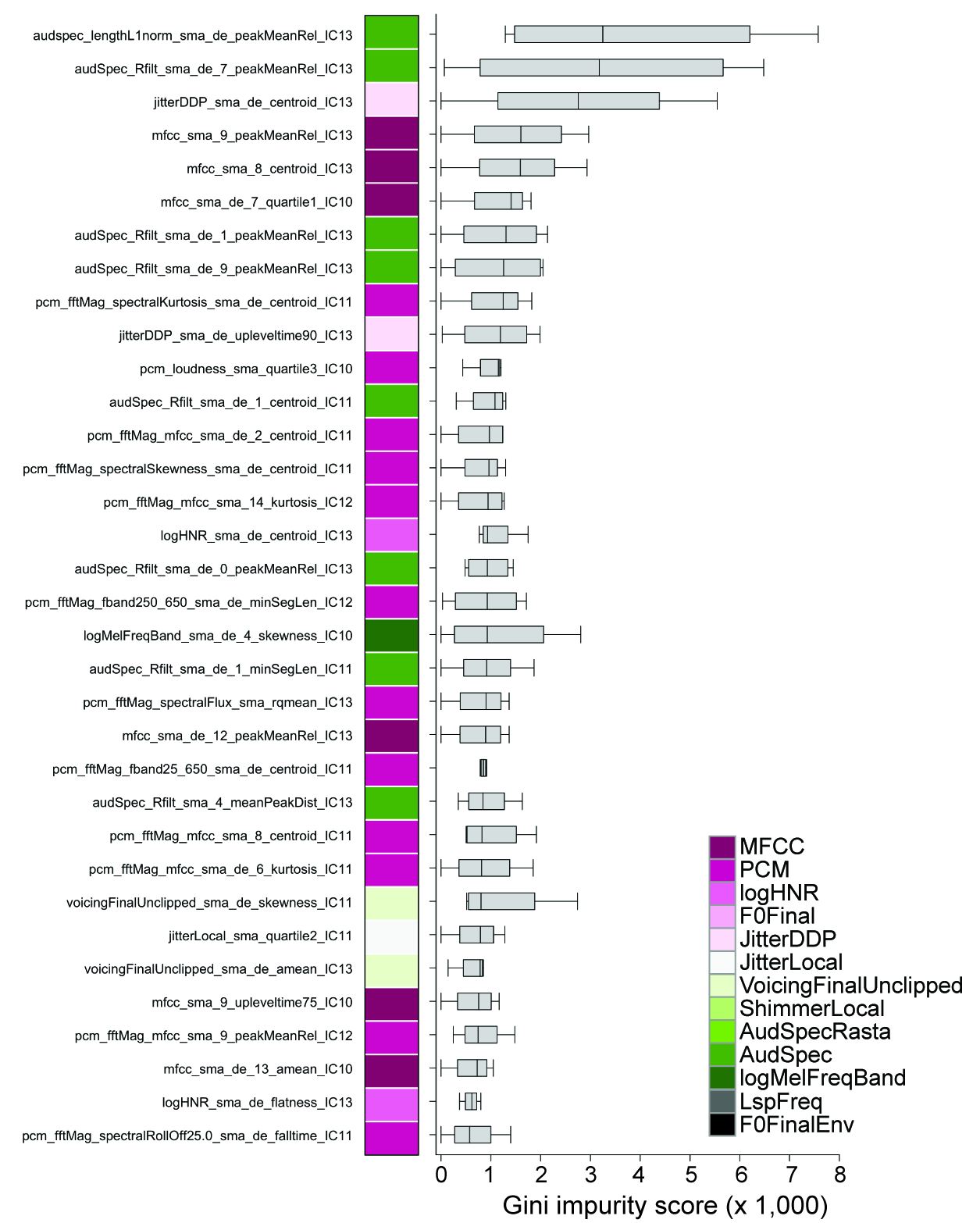

**Supplementary Figure 2.** Voice-features selected by Gini_c_. Voice-features with high Gini_c_ (Gini_c_ > 0.5) were selected as voice biomarkers. Gini impurity scores were measured from 1,000 repeated random stratified subsampling, score distributions were shown. Last four characters of voice-features (IC10, IC11, IC12, and IC13) indicated the origin of pre-defined feature set which OpenSmile provided.

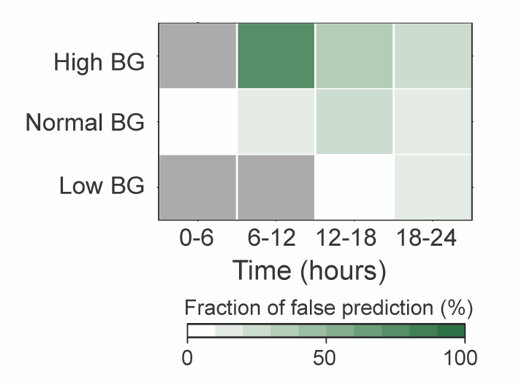

**Supplementary Figure 3.** Performance of blood glucose level prediction depending on time.

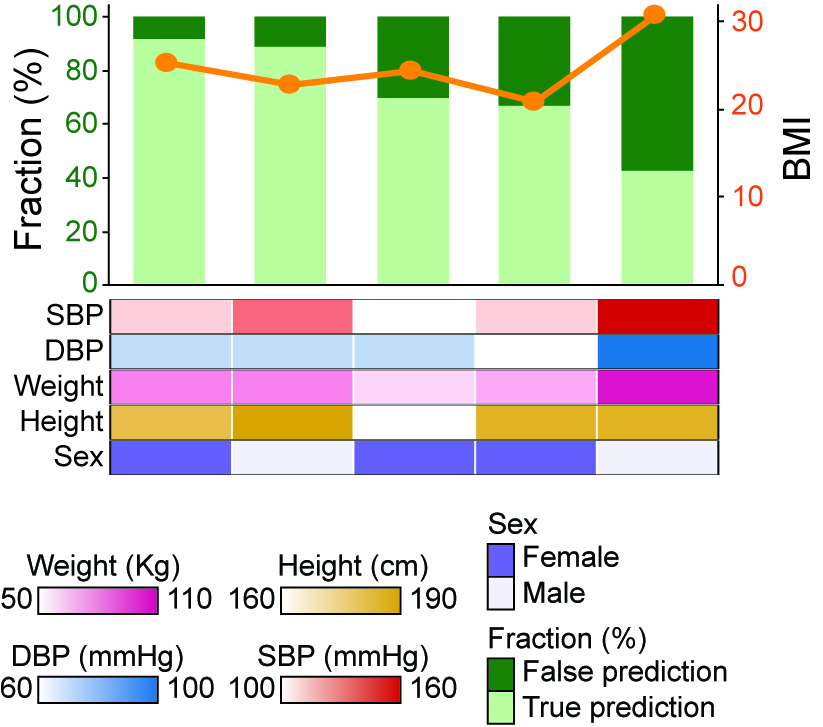

**Supplementary Figure 4.** Performance of blood glucose level prediction in the test set. Fractions of true (light green) and false (dark green) prediction depending on each individual were shown. SBP and DBP indicated systolic blood pressure and diastolic blood pressure, respectively.
